# *muat*: portable transformer-based method for tumour classification and representation learning from somatic variants

**DOI:** 10.64898/2026.04.01.715762

**Authors:** Prima Sanjaya, Esa Pitkänen

**Affiliations:** Institute for Molecular Medicine Finland (FIMM), HiLIFE, University of Helsinki, Finland; Applied tumour Genomics Research Program, Faculty of Medicine, University of Helsinki, Finland; iCAN Digital Precision Cancer Medicine Flagship, Finland

## Abstract

**Motivation:** Deep neural networks have proven effective in classifying tumour types using next-generation sequencing data. However, developing transferable models that work across heterogeneous operating environments remains challenging due to differences in cohort compositions and data generation protocols, privacy concerns, and limited computational capabilities.

**Results:** We introduce *muat*, a transformer-based software for tumour classification using somatic variant data from whole-genome (WGS) and whole-exome sequencing (WES). Building on previously developed MuAt and MuAt2 models, we distribute the software via Docker containers and Bioconda for deployment in high-performance computing (HPC) systems and Secure Processing Environments (SPEs). Using a downloadable MuAt checkpoint, we reproduce the performance reported in the original study on whole genome (PCAWG; 89% accuracy in histological tumour typing) and exome sequencing data (TCGA; 64% accuracy). Cross-cohort evaluation in Genomics England SPE achieved 81% accuracy without retraining and 89% following fine-tuning. As a demonstration of the software’s adaptability, we also deployed *muat* within the iCAN Digital Precision Cancer Medicine Flagship’s SPE and integrated it into a Nextflow-managed workflow.

**Availability and implementation:** *muat* is available through conda (www.anaconda.org/bioconda/muat) and GitHub (https://github.com/primasanjaya/muat), under the Apache 2.0 License.

**Contact:**; website: mlbiomed.net

## 1 Introduction

Accurate identification of tumour type and subtype is central to cancer diagnosis and treatment planning [1, 2, 3, 4]. While histopathological assessment remains the clinical standard, diagnostic uncertainty may arise in rare entities, poorly differentiated tumours, or limited biopsy material [5]. Molecular profiling provides complementary information and can improve diagnostic precision.

Recent studies have shown that deep neural networks trained on somatic mutation data can learn tumour-specific patterns directly from next-generation sequencing data [6, 7]. However, translating such models into routine research workflows is challenging. Genomic datasets are frequently hosted within Secure Processing Environments (SPEs), where internet access and software installation are restricted. Furthermore, inconsistencies in preprocessing pipelines, genome builds, and hyperparameter settings can compromise reproducibility when models are transferred across computing platforms. As a result, even publicly available research code is often difficult to deploy reliably within regulated genomic infrastructures.

To address these challenges, we developed *muat*, a portable implementation of mutation-level transformer models designed for reproducible execution across heterogeneous and secure computational environments. The framework formalises preprocessing, checkpoint packaging, and deployment procedures to enable deterministic training and inference across high-performance computing clusters and regulated data environments.

## 2 Implementation

*muat* is implemented in Python using PyTorch and provides a command-line interface supporting data preprocessing, model training, transfer learning, extraction of sample features, and inference (Fig. 1a). *muat* software implements the MuAt [7] and MuAt2 [8] transformer architectures as described in the original publications. The results reported in those studies were generated using the same core implementation that is now formalised and distributed as *muat*.

**Figure 1.**
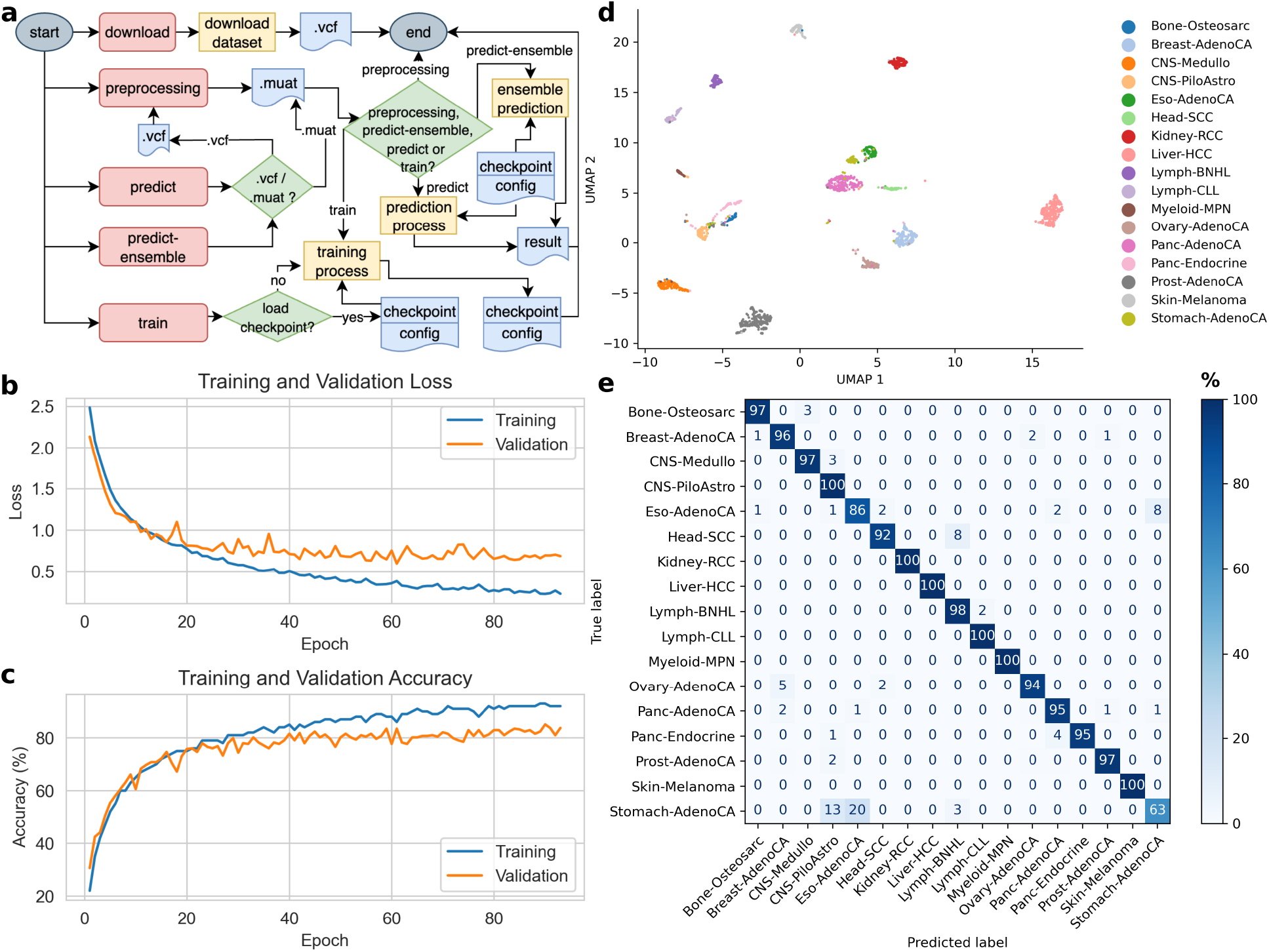
(**a**) Command-line workflow illustrating dataset download, preprocessing of VCF files into tokenised `.muat.tsv` format, training, checkpoint loading, and prediction. (**b-e**) Performance of a model trained on somatic SNVs and MNVs from publicly available whole-genome sequencing data (PCAWG). (**b**) Training and validation loss across epochs. (**c**) Training and validation accuracy across epochs. (**d**) UMAP projection of tumour-level representations learned by a MuAt model, showing clustering of tumour types in the latent embedding space. (**e**) Row-normalised confusion matrix (%) on the validation set, with rows showing true tumour types, columns predicted tumour types, and each row summing to 100%.

During preprocessing, somatic variants are converted into structured mutation tokens encoding the trinucleotide context, genomic position, gene and exon annotations, and coding strand orientation. The preprocessing pipeline supports somatic single-nucleotide variants (SNVs), multi-nucleotide variants (MNVs), short insertions and deletions (indels), structural variants (SVs), and mobile element insertions (MEIs), with positional and functional annotation. WGS and WES data in GRCh37/hg19 and GRCh38/hg38 reference assemblies are supported. To ensure compatibility with pretrained models trained on hg19 coordinates, *muat* converts data internally from hg38 to hg19. Preprocessing outputs are stored as reusable .muat.tsv files, which ensure that subsequent training or inference steps operate on identical token representations.

A central design feature of *muat* is reproducible checkpoint packaging. Each checkpoint stores the trained model weights together with the full preprocessing configuration, genome build information, and training hyperparameters required to reproduce the model state. Checkpoints follow a Hugging Face –compatible format (`huggingface.co`), using safe serialisation practices for portability and integrity across environments. The complete configuration is serialised as a structured JSON object and compiled into a single checkpoint file together the model parameters. By embedding architectural and preprocessing configuration within the checkpoint, *muat* ensures deterministic restoration across computational environments, and improves transparency by allowing the complete model configuration, preprocessing pipeline, and training context to be inspected directly from the checkpoint.

The software is distributed via Bioconda and as Docker containers through BioContainers for execution within SPEs. Pretrained checkpoints are included in the container image for easy inference without additional configuration.

## 3 Results

### 3.1 Reproducible benchmarking on publicly accessible data

To evaluate reproducibility and accessibility of the software independently of controlled-access datasets, we retrained MuAt using 1,901 publicly available ICGC whole-genome samples spanning 17 tumour types [9]. All data were obtained through the automated download functionality implemented in *muat* (Supplementary Data).

Using a stratified 9:1 train–validation split by tumour type, a model trained on SNVs and MNVs with positional and gene/exon/strand annotation achieved 85% validation accuracy (Fig. 1b,c). *muat* learned tumour-level representations formed well-separated clusters in latent space across tumour types (Fig. 1d). Misclassification patterns were largely confined to biologically related entities, with high per-class accuracy for most tumour types (Fig. 1e). This performance is comparable to that reported in the original MuAt study [7].

### 3.2 Reproducibility of published MuAt models

The software package *muat* provides pretrained MuAt checkpoints corresponding to the models described in the original study (Table 1) [7], where performance was reported using cross-validation on controlled-access PCAWG whole-genome and TCGA whole-exome datasets. PCAWG (WGS) checkpoints are organised by different mutation-type configurations, including SNV, SNV+MNV, SNV+MNV+indel, and SNV+MNV+indel+SV/MEI, with additional feature variants (motif-only, motif+positional, and motif+positional+annotation). For each input configuration, the best-performing model from each of the ten cross-validation folds is provided. For TCGA (WES), the same structure is available without SV/MEI-based models.

**Table 1:** Summary of pretrained MuAt checkpoints and reproduced performance.

| Dataset | Data type | Samples | Tumour types | Accuracy (%) |
| --- | --- | --- | --- | --- |
| PCAWG | WGS | 2,587 | 24 | 89 |
| TCGA | WES | 7,352 | 20 | 64 |

Inference can be performed using a single model via `–predict` or using an ensemble of models via `–predict-ensemble`. By specifying the mutation-type configuration, *muat* automatically selects the corresponding checkpoint. For ensemble prediction, *muat* aggregates outputs across the ten fold-specific models within the selected mutation-type configuration to produce a final prediction.

To enable reproducible deployment within a unified framework, these checkpoints were converted to the *muat* format while preserving the original model parameters. Using the standardised *muat* workflow on the CSC Puhti HPC cluster, here we re-evaluated these models across the ten fold models, obtaining a mean cross-validated accuracy of 89% on 2,587 PCAWG samples and 64% on 7,352 TCGA samples, consistent with the published results. Input configurations match the original study, with SNVs and MNVs (positional encoding) for PCAWG, and SNVs, MNVs, and indels (with positional and gene/exon/strand annotation) for TCGA. All checkpoints are available at https://huggingface.co/primasanjaya/muat-checkpoint/tree/main. These results confirm that the distributed checkpoints reproduce the performance of the original models.

### 3.3 Cross-environment portability

In the original MuAt study [7], PCAWG whole-genome models were trained on the CSC Puhti HPC cluster and subsequently evaluated within the Genomics England (GEL) SPE [10]. In that setting, the PCAWG-pretrained model achieved 81% classification accuracy on 9,796 tumour genomes spanning seven tumour types. These experiments were conducted using the same core implementation that is now included in the *muat* software, with mutation inputs adapted to the target environment, using SNVs, indels, and SVs in GEL.

Similarly, in the MuAt2 study [8], fine-tuning within the GEL SPE was performed using 14,527 whole-genome samples spanning 15 tumour types and 68 subtypes, using SNVs, indels, and SVs as input. Deep fine-tuning of the full model achieved 89% tumour type accuracy and 62% subtype accuracy. This training and transfer learning were likewise conducted using the implementation released in *muat*. In accordance with data governance policies, the fine-tuned checkpoint remains within the GEL SPE. The released version of *muat* corresponds to the implementation used in those studies.

To assess practical deployability in real-world research infrastructure, we integrated the *muat* Docker image within the iCAN SPE (`ican.fi`) into a Nextflow-managed workflow for containerised inference (see Supplementary Data). Execution required no modification of the container image or embedded checkpoint configuration. This demonstrates that *muat* operates as a modular component within workflow-orchestrated genomic analysis pipelines in regulated research environments.

## 4 Discussion

*muat* establishes a reproducible framework for mutation-level transformer models to be deployed in regulated genomic research infrastructures. By embedding preprocessing configuration, genome build compatibility, and architectural parameters directly within serialised checkpoints, model behaviour will be deterministic and reproducible across heterogeneous computational environments.

The distributed MuAt checkpoints correspond to the models described in the original publications [7], and their reported performance can be reproduced using the standardised *muat* workflow. MuAt2 checkpoints are maintained within the GEL SPE [8] and are not publicly distributed, but are available for reuse within the GEL SPE by authorised users. Deployment within secure environments, including the GEL and iCAN SPEs, further demonstrates that inference and transfer learning can be conducted without modification of container images or export of genomic data. Through versioned checkpoint packaging and container-based distribution, *muat* positions mutation-level transformer models as operational components within bioinformatics workflows, beyond typical research-only implementations.

While pretrained models can be imported into SPEs, cross-institutional training remains constrained by privacy and governance requirements. Federated or swarm learning strategies may offer future directions for collaborative model development without sharing raw genomic data [11, 12].

## 5 Conclusion

*muat* provides a reproducible and portable implementation of transformer-based tumour classification models with somatic variant data, supporting training, transfer learning and deployment in SPEs.

## 6 Data Access

This study analysed deidentified, publicly available data obtained from ICGC and TCGA data portals. Sequencing of human subjects’ tissue was performed by ICGC and TCGA consortium members under approval of local Institutional Review Boards (IRBs). Written informed consent was obtained from all human participants.

Data from the National Genomic Research Library (NGRL) used in this research are available within the secure Genomics England Research Environment. Access to NGRL data is restricted to adhere to consent requirements and protect participant privacy. Data used in this research include: somatic aggregated variant call (somAgg), somatic SV, and clinical metadata as provided through the LabKey in Genomics England Research Environment. The overview of genomics data used in the research, release v.16 (13th October 2022) can be found at https://re-docs.genomicsengland.co.uk/release16/. Preprocessed data is available in the Research Environment at /re_gecip/machine_learning/muat/data.

Access to NGRL data is provided to approved researchers who are members of the Genomics England Research Network, subject to institutional access agreements and research project approval under participant-led governance. For more information on data access, visit: https://www.genomicsengland.co.uk/research

## 7 Acknowledgements

We thank Nora Schreiber and Daniyar Karabayev for technical assistance. We also thank Anna Kuosmanen, Imre Västrik, Tua Karling, and Veronika Suni from the iCAN data team for their support. We acknowledge funding from the Research Council of Finland (#322675), the Sigrid Jusélius Foundation, and the Cancer Foundation Finland. We gratefully acknowledge the participants of the National Genomic Research Library (NGRL), whose contributions made this research possible. Secure access to the NGRL under project ID 837 was provided by Genomics England, which delivers the NGRL in partnership with NHS England, and is wholly owned by the UK Department of Health and Social Care. The NGRL contains participants’ health data collected by the NHS as part of their care, along with samples and data from their participation in research, for which fully informed consent has been obtained. This includes genomic and clinical data provided through the NHS Genomic Medicine Service, as well as data obtained through research studies, including the 100,000 Genomes Project and the Generation Study, both of which are delivered in partnership with the NHS, and from other research cohorts involving external collaborators. The authors wish to acknowledge CSC – IT Center for Science, Finland, and iCAN Digital Precision Cancer Medicine Flagship for computational resources.

## 8 Supplementary Data

Supplementary data are available at Bioinformatics online.

