## Supplementary Data for "*muat*: portable transformer-based method for tumour classification and representation learning from somatic variants"

### Downloading public data

To train and test the models, *muat* provides a convenience command (`download`) to download publicly available data from the International Cancer Genome Consortium (ICGC). ICGC hosts tumour whole-genome sequencing (WGS) data through an Object Storage Bucket (https://docs.icgc-argo.org/docs/data-access/icgc-25k-data). For example, the command in Supplementary Data 1 will download 1,901 WGS samples across 17 tumor types from the ICGC subset of the PCAWG.

```
muat download --pcawg
```

Supplementary Listing 1: Command for downloading publicly available PCAWG data.

### Preprocessing input data

Variant Calling Format (VCF) files of tumour somatic mutations—either gzipped or unzipped and based on the human genome reference GRCh37 (hg19) or GRCh38 (hg38)—can be used as direct input for prediction, in which case *muat* automatically performs all required preprocessing internally. For training, however, *muat* expects preprocessed and tokenised input files generated from these VCFs.

To run preprocessing manually, the `preprocess` command can be used to generate tokenised input. This is useful for inspecting the processed features or reusing the input across multiple runs. The command performs all necessary steps in a single execution: (1) extracting the 3-bp sequence motif, genomic positions in hg19 coordinates, and annotations indicating whether the mutation lies within a gene or exon, along with its orientation with respect to the transcript orientation, then (2) tokenising these features into unique indices using predefined dictionaries.

The result is a single output file with the extension *.muat.tsv*, which can be provided as input using the `--no-preprocess` flag to skip internal preprocessing. Below are examples of how to preprocess the data:

```
#example for hg19 VCF
muat preprocess --vcf --hg19 <hg19 path> --input-filepath <vcf path>
#example for hg38 VCF
muat preprocess --vcf --hg38 <hg38 path> --input-filepath <vcf path>
```

Supplementary Listing 2: Command for preprocessing VCF files

### Training MuAt models from scratch

| **prep_path** | **class_name** |
|---|---|
| <absolute_path>/sample1.muat.tsv | Breast-AdenoCA |
| <absolute_path>/sample2.muat.tsv | Ovarian-AdenoCA |

Table 1: Format of the training data file used by muat (TSV).

To train MuAt models, first prepare a tab-separated values (TSV) file containing the absolute paths of the preprocessed files (specified in the **prep_path** column) and their corresponding labels (**class_name**) for both the training and validation splits (see Table 1). Next, define the hyperparameters for model training, such as the number of epochs, learning rate, number of MuAt encoder layers, and other relevant settings. Finally, specify the mutation types to use (*i.e.* `snv`, `snv+mnv`, `snv+mnv+indel`, `snv+mnv+indel+sv`, `snv+mnv+indel+sv+neg`) and whether to include motif information alone (*i.e.* `--use-motif`) or to incorporate position (i.e. `--use-position`) and annotation features (*i.e.* `--use-ges`). Upon training completion, a checkpoint is written which stores all configurations and model parameters for later inference or fine-tuning. Supplementary Data 3 is an example command for training a MuAt model from scratch:

```
muat train from-scratch --mutation-type <mutation type> --use-motif --
   use-position --use-ges --train-split-filepath <train split> --val-
   split-filepath <validation split> --save-dir <save dir> --epoch <
   epoch> --learning-rate <learning rate> --batch-size <batch size> --n
   -layer <n layer> --n-head <n head> --n-emb <n emb> --mutation-
   sampling-size <n mutation sampling>
```

Supplementary Listing 3: Command for training from scratch

### Training from pretrained models and finetuning

Fine-tuning an existing model on new data is also supported by *muat*. Previously, we fine-tuned the MuAt models to create MuAt2 models. For example, if a user has a pre-trained checkpoint, they can continue training it on a different dataset and labels.

To do this, the user needs to specify a new training and validation data split, including the labels, as described in the previous section. To include additional labels (*e.g.*, tumour subtypes in MuAt2), the user can add the **subclass_name** column to the TSV file. The user must also define the training hyperparameters, such as the number of epochs, learning rate, and batch size. However, the model hyperparameters cannot be adjusted because they are loaded from the checkpoint.

Supplementary Data 4 is an example command for training a MuAt model from a pre-trained checkpoint:

```
muat train from-checkpoint --ckpt-filepath <checkpoint path> --mutation
   -type <mutation type> --train-split-filepath <new train split> --val
   -split-filepath <new validation split> --save-dir <save dir> --epoch
    <epoch> --learning-rate <learning rate> --batch-size <batch size>
```

Supplementary Listing 4: Command for training from checkpoint

### Prediction

Predictions (*i.e.*, classifying tumour types) can be made directly from raw VCF files, which are automatically preprocessed by *muat*. Alternatively, users may use pre-tokenised input using the `--no-preprocess` flag, in which case the input must already be processed using `muat preprocess`.

To run model inference, users must choose whether to load a pre-trained model from a checkpoint or specify the mutation type for the input file. To simplify the prediction process, MuAt pre-trained models [?] are included in the software for users who have not had any prior training and wish to make predictions.

MuAt supports both whole-genome sequencing (WGS) and whole-exome sequencing (WES) data. The subcommand `predict wgs` should be used for WGS input, while `predict wes` is used for WES data, as the underlying models may differ.

To run a quick inference on WGS data (replace `--hg19` with `--hg38` if the VCF was generated using GRCh38), execute command in Supplementary Data 5:

```
muat predict wgs --hg19 <hg19 path> --mutation-type <mutation type> --
    input-filepath <vcf path> --result-dir <result dir>
```

Supplementary Listing 5: Command for prediction from VCF file

This command will preprocess the VCF file and generate predictions using the MuAt pre-trained model for the specified mutation type. Alternatively, users can run predictions directly on preprocessed files and specify a pretrained model from a checkpoint (Supplementary Data 6):

```
muat predict wgs --no-preprocess --ckpt-filepath <checkpoint path> --
    input-filepath <preprocessed file> --result-dir <result dir>
```

Supplementary Listing 6: Command for prediction from preprocessed file

Tumour class probabilities and predicted labels for each sample will be written to the specified result directory.

### Benchmarking

To reproduce the best results from MuAt [?], the best approach is to ensemble models from 10-fold cross-validation. This software provides the best ensemble models for both WGS and WES data. To run the benchmark MuAt models, use the commands in Supplementary Data 7 (replace `muat-wgs` with `muat-wes` for WES data benchmarking):

```
muat predict-ensemble muat-wgs --hg19 <hg19 path> --mutation-type <
    mutation type> --input-filepath <vcf path> --result-dir <result dir>
```

Supplementary Listing 7: Command for prediction using muat benchmark models

This will download the ensemble checkpoints from https://huggingface.co/primasanjaya/muat-checkpoint and generate predictions by averaging probabilities over 10 ensemble models.

### Using *muat* via a container

Prebuilt container images for *muat* are available from BioContainers or can be built directly from source code. As an example, we below show how to run containerized *muat* in iCAN-DP SPE, which is based on Azure Cloud. The container must first be pulled from BioContainers or built locally, and then uploaded to the iCAN **public** Azure Container Registry (ACR). Once uploaded, it can be pulled from the iCAN **private** ACR within the iCAN-DP SPE.

The steps in Supplementary Data 8 outline the process for transferring muat from BioContainers or local to the iCAN **public** ACR:

```
REGISTRY_NAME="<ican_public_acr>"
MUAT_VERSION="<tag>" # Update version if needed
# Step 1: Log in to Azure
# Step 2: Log in to the ican public acr
# Step 3: Pull muat from BioContainers or build docker from source code
docker pull quay.io/biocontainers/muat:"$MUAT_VERSION"
# Step 4: Tag the image for the ican public acr
docker tag quay.io/biocontainers/muat:"$MUAT_VERSION" "$REGISTRY_NAME".
    azurecr.io/muat:"$MUAT_VERSION"
# Step 5: Push the image to the ican public acr
docker push "$REGISTRY_NAME".azurecr.io/muat:"$MUAT_VERSION"
```

Supplementary Listing 8: Steps for pushing docker to iCAN public ACR

Details on how to login to iCAN ACR and are documented https://ican-1.gitbook.io/handbook/software-and-tools/. Once the image is uploaded to the iCAN **public** ACR, *muat* can be used within the iCAN-DP SPE by pulling the container from the iCAN **private** ACR and running the comman in Supplementary Data 9:

```
REGISTRY_NAME="<ican_private_acr>"
MUAT_VERSION="<tag>"  # Match the pushed version
# Step 1: Pull the muat image from the ican private acr
docker pull "$REGISTRY_NAME".azurecr.io/muat:"$MUAT_VERSION"
# Step 2: Run muat --help interactively
docker run -it "$REGISTRY_NAME".azurecr.io/muat:"$MUAT_VERSION" muat -h
```

Supplementary Listing 9: Steps for pullling docker to iCAN private ACR

In GEL SPE, *muat* can be imported in similar fashion via artifactory. Details on using GEL artifactory are written in `https://re-docs.genomicsengland.co.uk/hpc_containers/`.

### Pipeline integration

To demonstrate integration into analysis workflows, we provide an example of a minimal Nextflow pipeline that runs *muat* prediction on an input VCF file using a Docker container. This pipeline automates *muat* predictions: it pulls the Docker image, executes `muat predict` on input VCFs, and saves the results. This allows *muat* to be easily incorporated into reproducible pipelines in SPEs or larger analysis pipelines.

Supplementary Data 10 shows the core Nextflow process that calls *muat* within the pipeline:

The full pipeline script (`main.nf`) and associated `nextflow.config` file defines default values, including the mutation type (`snv+mnv`), paths to the input VCF and reference genome, and compute resources are available in the muat github repository.

Finally, to execute the pipeline, use the command in Supplementary Data 11:

```
process runMuatPredict {
    input:
    path input_vcf
    path reference
    val result_dir

    output:
    path result_dir

    script:
    """
    muat predict wgs \
        --hg19 ${reference} \
        --mutation-type '${params.mutation_type}' \
        --input-filepath ${input_vcf} \
        --result-dir ${result_dir}
    """
}
```

Supplementary Listing 10: example script of main.nf

```
nextflow run main.nf
--input_vcf <vcf path>
--reference <hg19 or hg38 path>
--result_dir <result dir>
-profile docker
```

Supplementary Listing 11: Command for running nextflow for muat prediction
